# Effect of environmental conditions on the wing morphometric variation in *Aedes aegypti* (Diptera: Culicidae) in India

**DOI:** 10.1101/2024.02.01.578359

**Authors:** Gaurav Sharma, Rakesh Bhutiani, Devojit Kumar Sarma

## Abstract

*Aedes aegypti* an efficient vector of different arboviral diseases, is a major concern for public health globally. Originating from Africa, this vector has now invaded most of the parts of the world, which shows its thriving adaptability against diverse ecological conditions. In India too, *Ae. aegypti* has been found almost everywhere and along with *Ae. albopictus*, it contributed more than 0.2 million cases of dengue in the year 2022. Due to heterogeneous environmental settings in India, this vector has shown various intraspecific variations including its behavioral, genetic, and physiological characteristics. Thus, the present study hypothesized that there will be some differences in wing morphometrics across the country for this species. Considering this, we have sampled adults and immatures of *Ae. aegypti* from 11 distinct locations, representing six varied climatic regions of India. The immatures were reared to adult and the right wing was used to score the morphometric variations. A significant variation in wing size and shape was observed. The Himalayan region (Srinagar CS: 1.92 ± 0.08 mm) supports the shortest (CS: 2.04 ± 0.09 mm) wing size mosquitoes while the Deccan Peninsular region (Nagpur CS: 4.48 ± 0.06 mm) exhibits the largest (CS: 4.46 ± 0.10 mm) wing size mosquito. After excluding the allometric effect, the semi-desert region (Kota) showed the greatest wide variety of wing shapes corresponding to a larger morphospace in CVA analysis. In addition, region-wise wing size was found positively correlated with temperature (64%) and altitude (82%) but no directional trend was detected for wing shape characteristics. Conclusively, the study suggests the existence of varied population structures of *Ae. aegypti* in India based on wing morphometric analysis. This finding will be helpful towards focused actions and early measures to reduce the impact of these diseases carrying mosquitoes on public health.

## Introduction

Dengue fever, the most prevalent mosquito-borne illness, has witnessed a substantial increase in reported cases, with a 10-fold surge observed from 2000 to 2019 globally (WHO, 2023). In India, more than 0.2 million cases have been reported in the year 2023 (NCVBDC, 2023). *Aedes aegypti* (Linnaeus), is the primary vector, responsible for this deadly disease. It is also the potential vector of several other epidemiologically important diseases including Chikungunya, and Zika in India (NCVBDC, 2023).

Historically, *Ae. aegypti* preferred non-human hosts and inhabited tropical forests, with larvae breeding in tree holes. However, as the human population expanded, *Ae. aegypti* population underwent evolutionary changes to adapt to human habitats. It is believed that this mosquito species has invaded other parts of the world through human movements and trades from Central Africa (Tabachnick, 1991). Consequently, diverse ecological and geographical conditions lead to changes in *Ae. aegypti* mosquito in context to their morphology, behavior, and genetics (Louise et al., 2015; Sumitha et al., 2023; Sharma et al., 2023).

Controlling the *Ae. aegypti* population is considered the best effective management strategy to prevent the expansion of vector-borne diseases, making an understanding of population dynamics important. Population dynamics studies related to behavior and genetics, provide vital information about how mosquitoes react in different environments (Barrera et al., 2011; Grech et al., 2015; De et al., 2022; Sharma et al., 2023; Sumitha et al., 2023).

Morphological analysis, particularly, Geometric morphometrics (GM) provides another important monitoring tool to understand the inter and intraspecific variations in Culicidae (Lorenz et al., 2017). This approach allows to investigate and comprehend morphological differences influenced by factors such as gender (Christe et al., 2016), geographical location (Hounkanrin et al., 2023), phylogenetic relationships (Lorenz et al., 2015), ecological associations (Carreira et al., 2011), impact of various treatments such as temperature (Aytekin et al., 2009), demographic factors (Wilk-da-Silva et al., 2018), altitude (Leyton Ramos et al., 2020) and landscape types (Hounkanrin et al., 2023). These studies have shown that wing size and shape are the reliable morphological traits used in geometric morphometrics.

Although most of the Indian cities experience a tropical to subtropical climate, there exists a local variation in topography, and land use contributing to micro-climatic variations influencing dengue vector abundance and epidemiology (Sarma et al., 2022). Due to this environmental heterogeneity in India, six different geo-climatic regions can be identified: Himalayan, Desert, Semi Desert, Deccan Peninsular, Coast, and North East (Dimitrova and Bora, 2020). It is evident that *Ae. aegypti* have invaded all these climatic regions, having different genetics (Sumitha et al., 2023) and behavioral differences (Sharma et al., 2023) in India, leading to the hypothesis that there will be a significant difference in the wing morphometrics. Thus, a collection of *Ae. aegypti* was conducted from 11 distinct locations in India, which correspond to six different climatic regions, and their wing morphometrics were studied to assess any difference in the wing morphometrics of this vector species.

## Material and Methods

### Study sites and sample populations

The *Ae. aegypti* mosquitoes were collected from a total of 11 locations (Jodhpur, Kota, Sri Ganganagar, Srinagar, Hoshiarpur, Nagpur, Raipur, Kolkata, Vishakhapatnam, Guwahati, and Itanagar) representing six different geo-climatic regions (Desert, Semi Desert, Himalaya, Deccan Peninsula, Coast and North East) in India (Figure 1). Detailed information about the sampling sites of each population is provided in Supplementary Table 1. Mosquitoes were sampled within human dwellings in localities.

**Figure 1.**
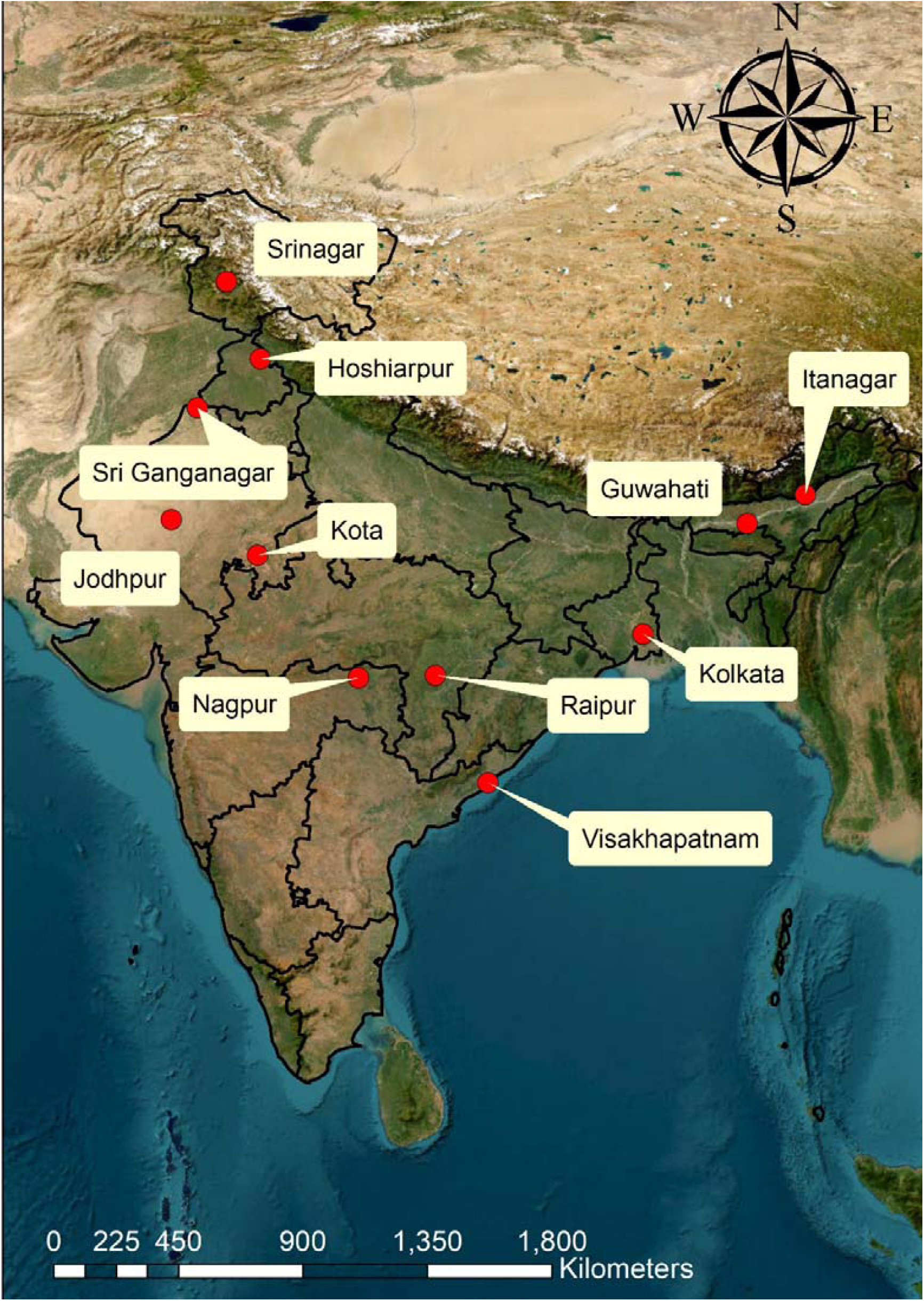
**Sampling location of *Aedes aegypti* in India**

### Sample Collection

Adult mosquitoes were collected by using a mouth aspirator whereas fourth instar larvae and pupae stages were collected using a glass pipette from most probable *Ae. aegypti* breeding habitats. Larvae and pupae were reared up to adult emergence. Field-collected and emerged adults were anesthetized using cotton soaked in 0.5ml of chloroform and identified morphologically using standard taxonomic keys (Tyagi et al., 2015). All adult specimens were kept under -20 °C for morphometric study.

### Wing preparation

The right wing of female *Ae. aegypti* mosquito was segregated from the thorax and mounted over a microscope slide (15 mm x 15 mm) with a coverslip. Each wing was then photographed under 40x magnification using ImageView software (version x64, 4.11.18012.20201123) with STEMI 305 stereomicroscope.

### Landmark digitisation

A total of 18 landmarks were digitized using ImageJ ver. 1.54d (Schneider et al., 2012) software following methods described by Hounkanrin et al. (2023). The wing picture and digitization of landmarks were done by one author (GS) and repeated three times to minimize error. Total 239 wing specimens of female *Ae. aegypti* (Jodhpur - 22, Kota - 8, Sri Ganganagar - 27, Srinagar - 13, Hoshiarpur - 24, Nagpur - 27, Raipur - 13, Kolkata - 27, Vishakhapatnam - 20, Guwahati - 24, and Itanagar – 32) were used. The representation of 18 landmarks and their coordinates are provided in Figure 2 and Supplementary File 1 respectively.

**Figure 2.**
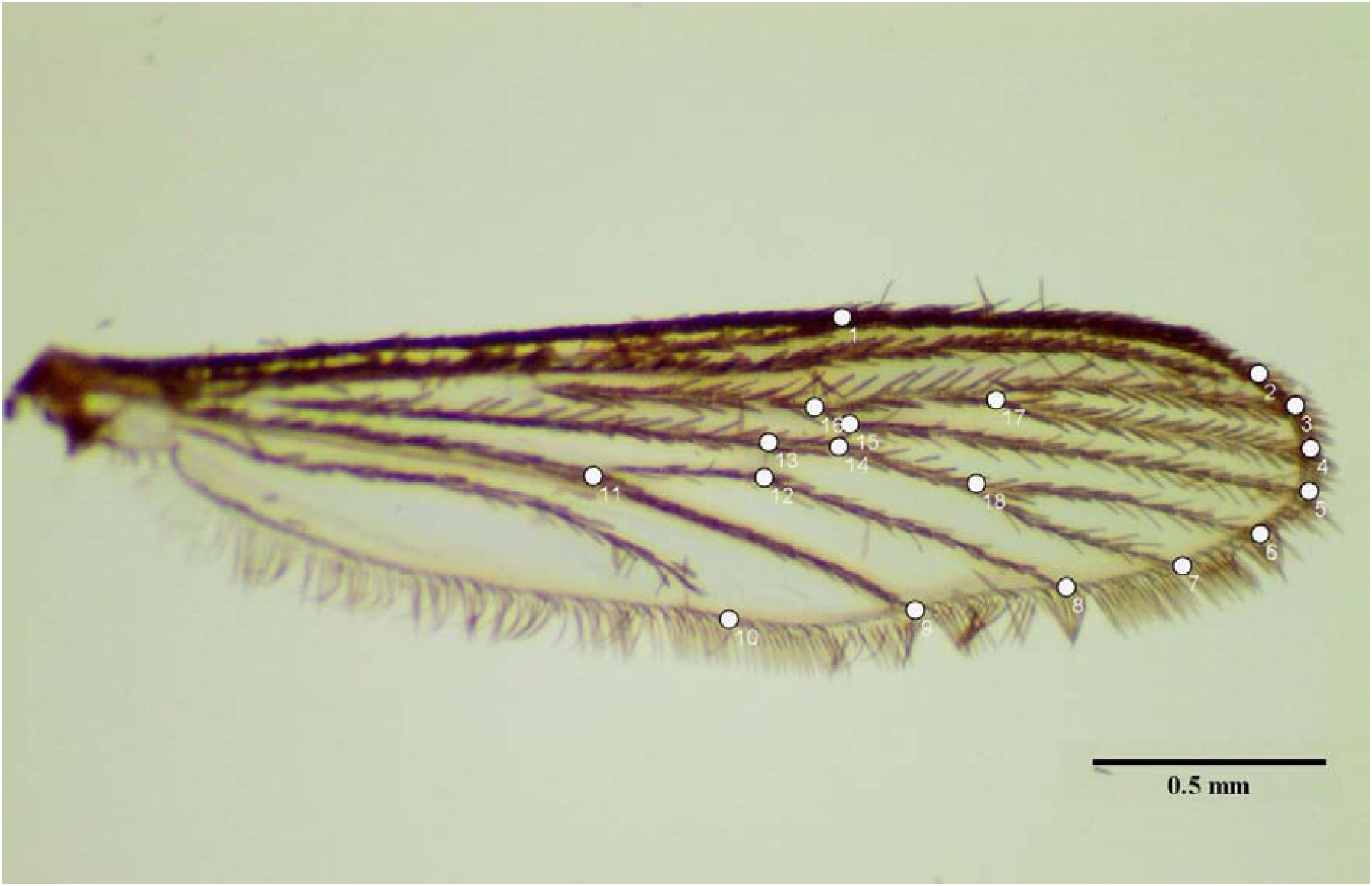
**Position of the 18 landmarks (digitized from 1st to 18th respectively) on a female *Aedes aegypti* right wing.**

### Wing size and shape estimation

The centroid size (CS) representing wing size, was calculated from raw wing coordinates data. Centroid size is defined as the square root of the total squared distances measured from the centroid to each landmark, and it can be employed as a proxy for wing size (Dujardin, 2008). Coordinates were imported into the Python Environment (Bisong, 2019) and processed with the help of pandas and the Numpy library. These coordinates were aligned by performing Procrustes superimposition (Figure 3) to visualize the position of each landmark for each specimen using MorphoJ ver. 1.08.01 (Klingenberg, 2011). The mean positions of the landmarks per population were calculated using the scipy.spatial library and visualized using matplotlib.pyplot (Figure 4). To statistically compare the mean CS among specimens from different populations and regions, ANOVA (multiple sample comparison test) was performed using STATGRAPHICS Plus version 5.0.

**Figure 3.**
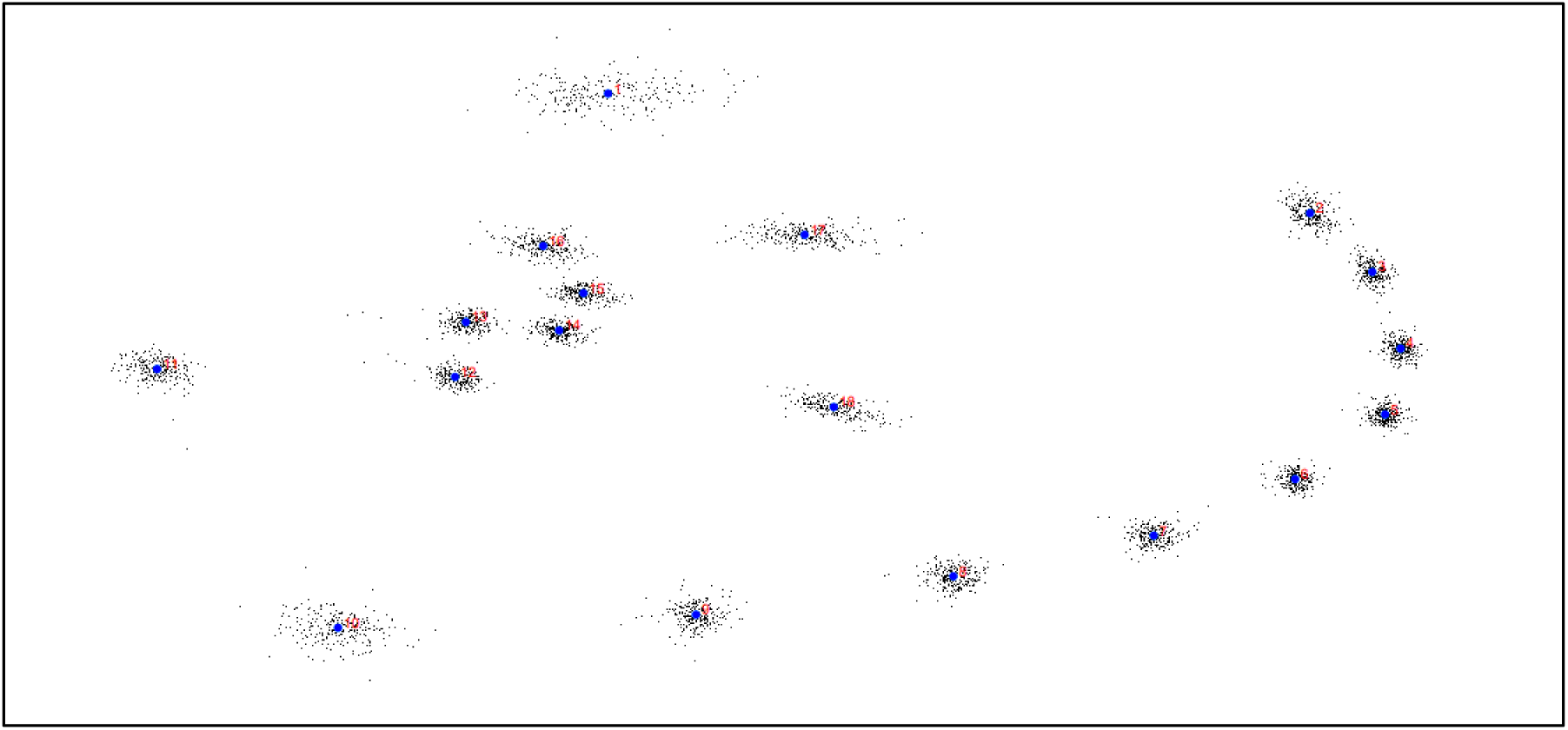
**Landmark positions derived from Procrustes superimposition.**

**Figure 4.**
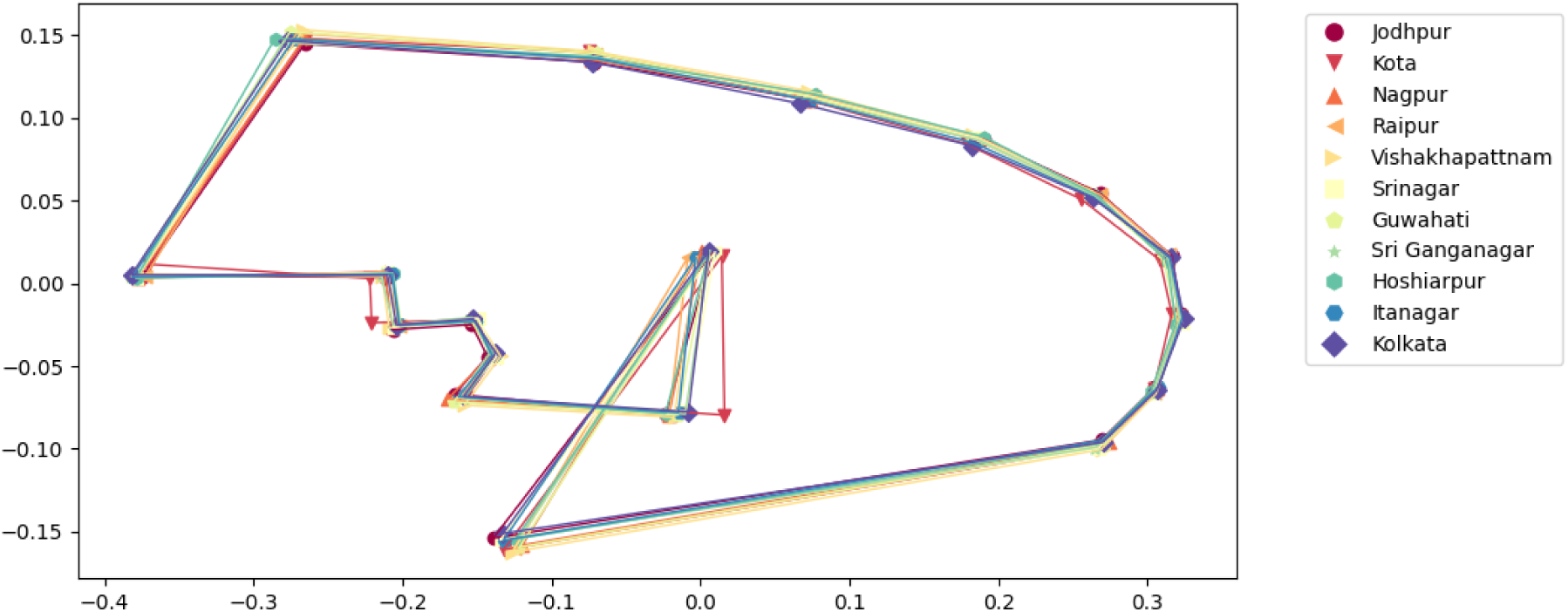
**Mean position of the 18 landmarks by different populations of India.**

The assessment of wing shape involved the importation of wing coordinates in a text-delimited format using MorphoJ ver. 1.08.01 (Klingenberg, 2011). To conduct a comparative analysis, Procrustes ANOVA was performed utilizing the same software. Following this, Canonical Variate Analysis (CVA) was applied to visually represent shape variations based on Procrustes coordinates. Additionally, Mahalanobis distances were computed using MorphoJ ver. 1.08.01 software to further quantify the differences in wing shape.

To ensure the species confirmation, all the specimens were analyzed morphologically as well as at the molecular level following the method described by Higa et al. (2010) targeting the ITS region of the genus *Aedes*. Both populations and regions were considered for the comparative analysis of wing size and wing shape.

### Phenetic relationship

Neighbour-Joining (NJ) tree was constructed with 100 bootstrap replicates based on Mahalanobis distances obtained through pairwise comparison of wing specimens via CVA using PAST software v.4.03 (Hammer et al., 2001) to illustrate the phenetic relationships among the populations and climatic regions.

### Allometric effect

To understand the influence of wing size on wing shape (allometry), the multivariate regression of the Procrustes coordinates (dependent variable) against log CS (independent variable) was analyzed using a permutation test with 10,000 randomizations utilizing MorphoJ ver. 1.08.01.

### Environmental variables

Historical environmental data including temperature, relative humidity, and precipitation, spanning the years 1981 to 2022, were obtained from the NASA, POWER LARC Data Access Viewer (https://power.larc.nasa.gov/data-access-viewer/). To assess and compare these variables among different populations, a multiple-sample comparison ANOVA was conducted utilizing STATGRAPHICS Plus ver. 5.0.

### Correlation between CS and Environmental Variables

The multivariate correlation analysis of the centroid sizes and different environmental factors including temperature, relative humidity, precipitation, latitude, longitude, and altitude was performed using PAST v.4.03.

## Results

The present study conducted wing morphometric analyses using a total of 239 wing specimens obtained from 11 distinct populations of *Ae. aegypti* (Supplementary Table 1) in India. The mean CS estimated from the coordinates of 18 wing landmarks, exhibited a significant difference among the sampling populations (F ratio = 241.08, *p* = <0.0001). Nagpur population showed the highest CS (4.48 ± 0.06 mm) while Srinagar displayed the lowest CS (1.92 ± 0.08 mm) (Figure 5). Similarly, a significant difference was also observed among the sampling regions (F ratio = 98.56, *p* = <0.0001) for the mean CS. The Deccan Peninsula region showed the highest CS (4.46 ± 0.10 mm), while the Himalaya region had the lowest (2.04 ± 0.09 mm) (Figure 6).

**Figure 5.**
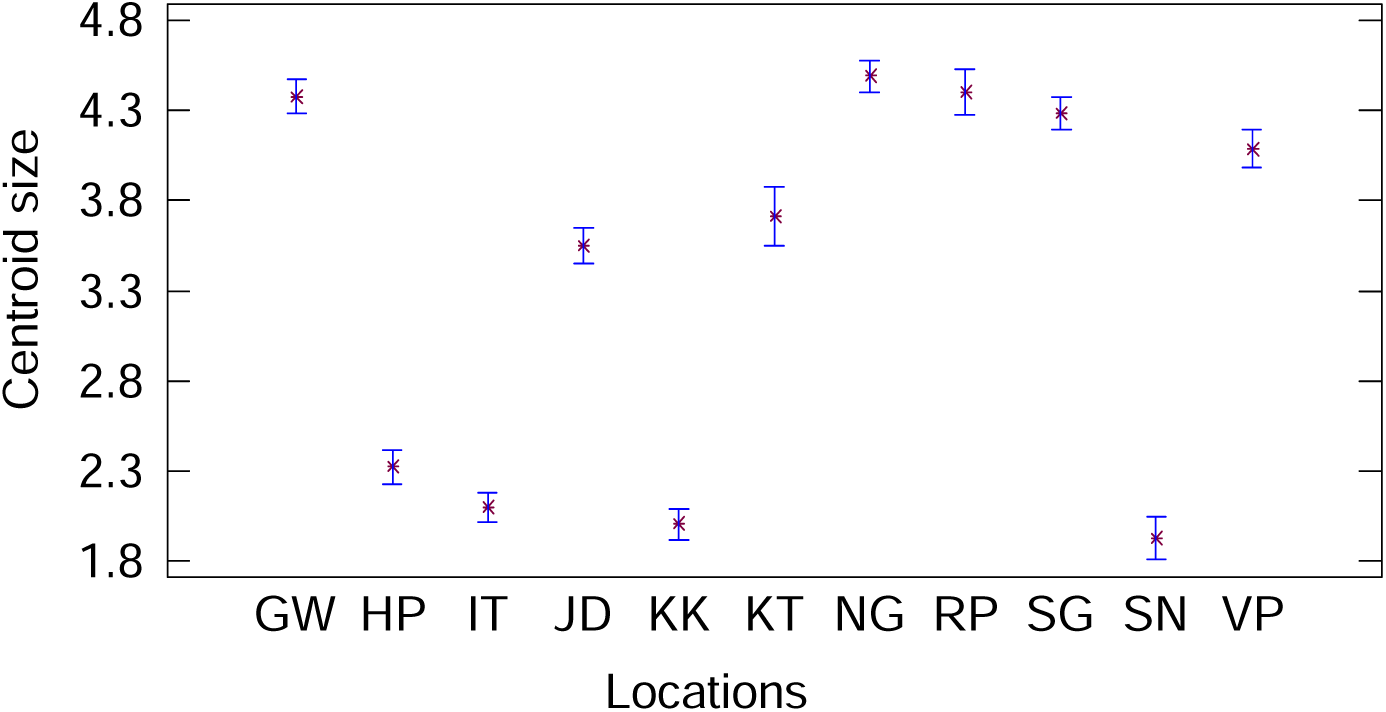
**Comparison of mean centroid size by populations (Abbreviation: GW – Guwahati, HP – Hoshiarpur, IT – Itanagar, JD – Jodhpur, KK – Kolkata, NG – Nagpur, RP – Raipur, SG – Sri Ganganagar, SN – Srinagar, VP - Visakhapatnam.**

**Figure 6.**
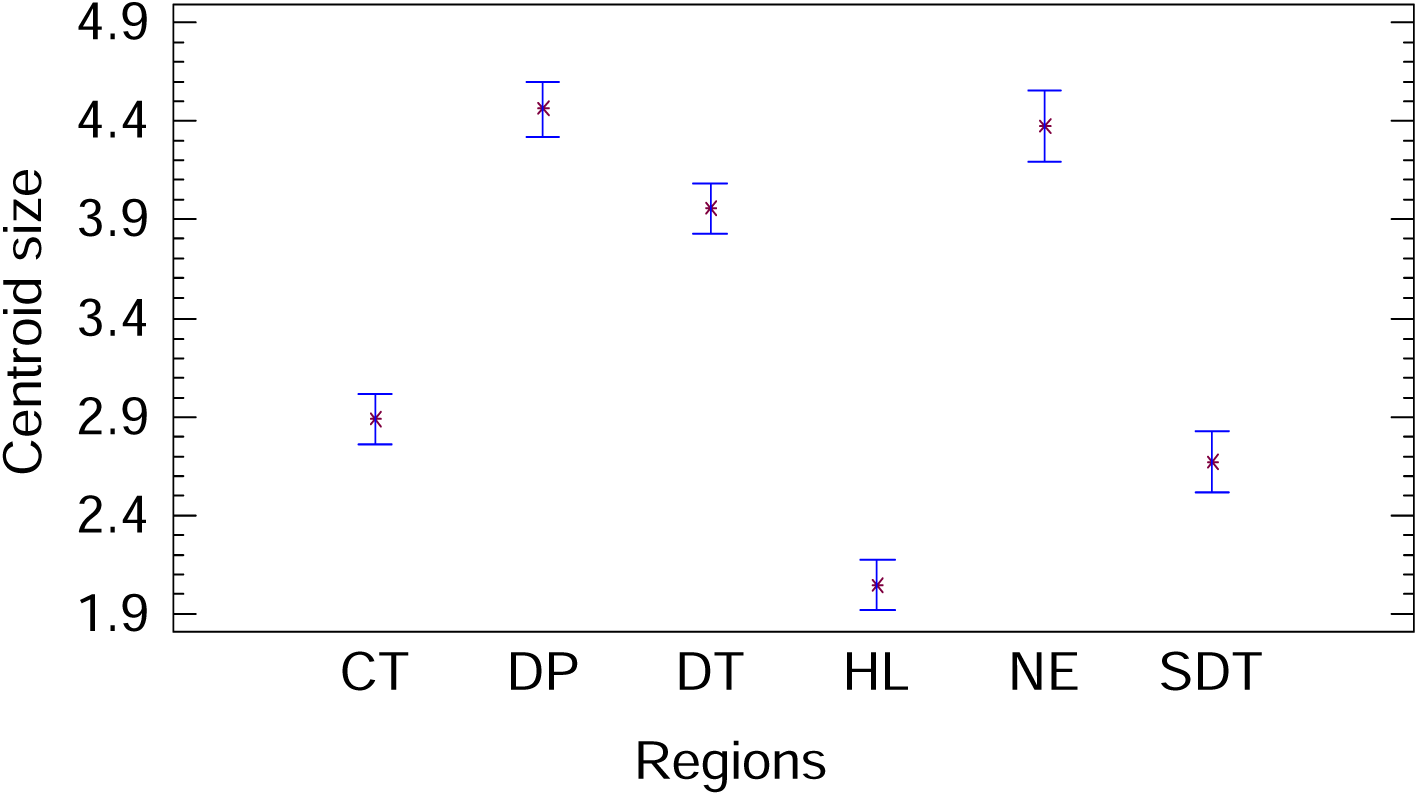
**Comparison of mean centroid size by regions (Abbreviation: CT – Coasts, DP – Deccan Peninsula, DT – Desert, HL – Himalaya, NE – North East, SDT – Semi-desert)**

In the analysis of wing shape variation, Procrustes ANOVA revealed significant differences both among the populations (F ratio = 2.71, *p* <0.0001) and regions (F ratio = 1.92, *p* < 0.0001). Canonical Variate Analysis (CVA) accounted for 38.89 % of the total variance on the first two canonical variates (CV1 = 22.21%, CV2 = 16.68%) among the populations (Figure 7). The scatter plots of CV1 and CV2 showed overlapping wing shape variations among the populations. At a region level (Figure 8), CVA explained 61.21% of the total variation (CV1 = 31.10%, CV2 = 28.11%), with an overlapping scatter plot.

**Figure 7.**
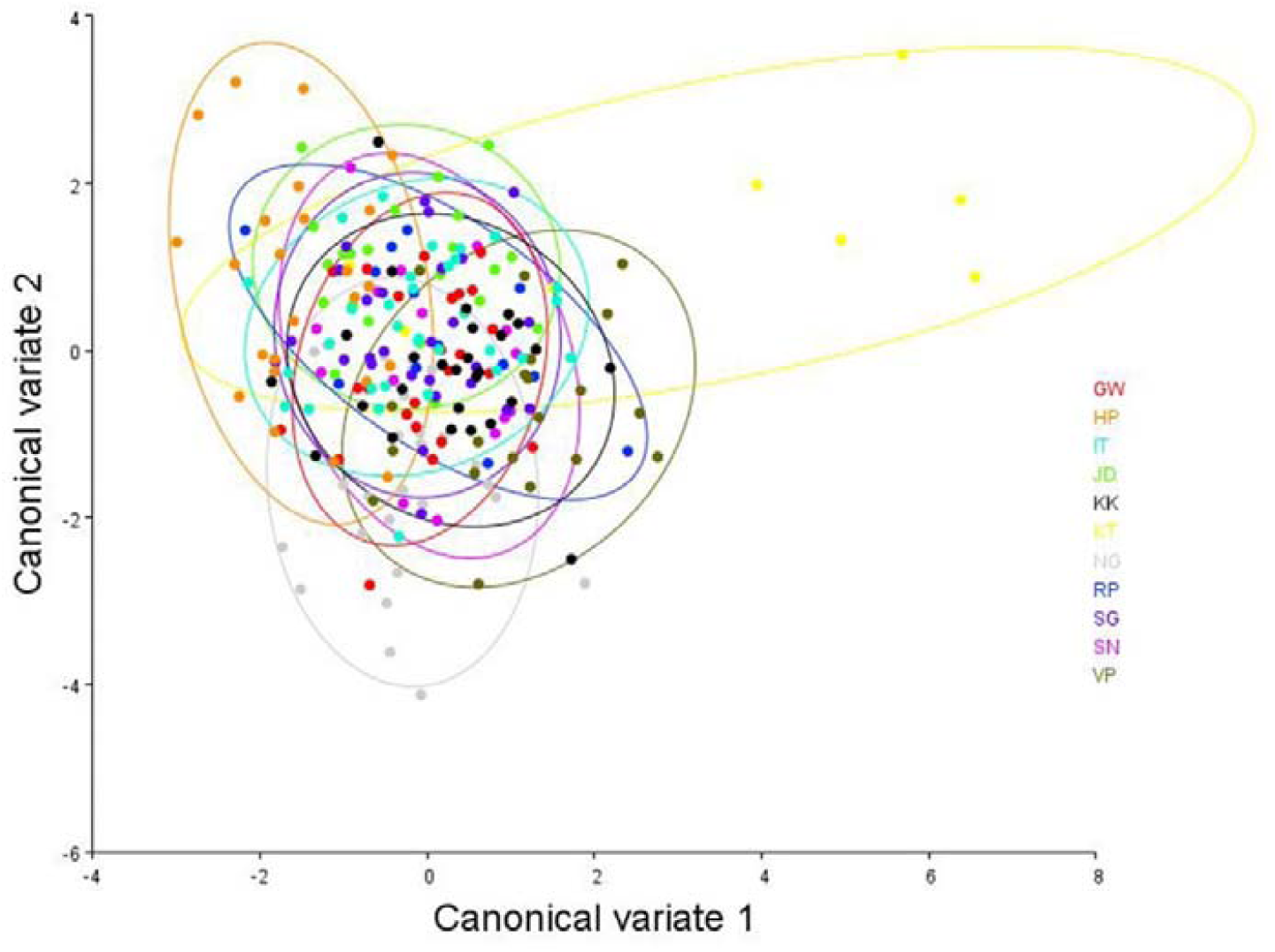
**Canonical Variate Analysis of the Procrustes coordinates of *Aedes aegypti* by populations. (Abbreviations: GW – Guwahati, Hp – Hoshiarpur, IT – Itanagar, JD –Jodhpur, KK – Kolkata, KT – Kota, NG – Nagpur, RP – Raipur, SG – Sri Ganganagar, SN – Srinagar, VP – Visakhapatnam)**

**Figure 8.**
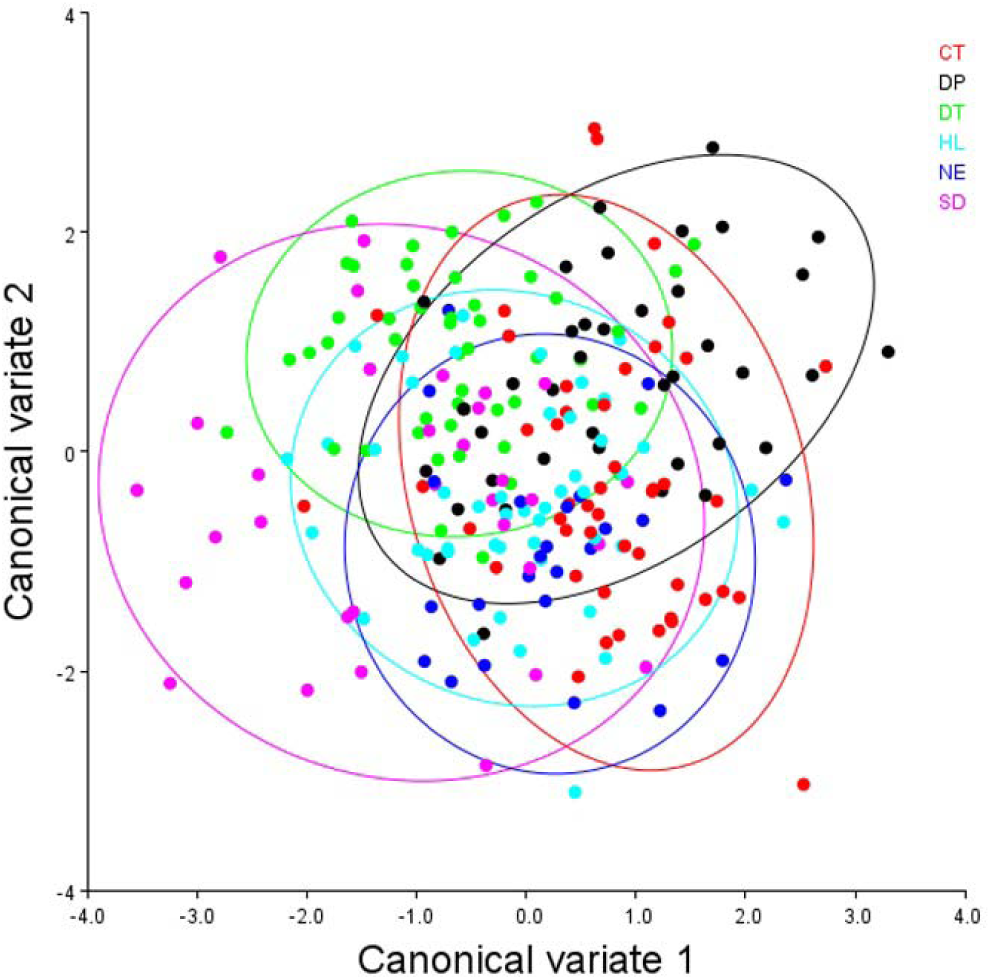
**Canonical Variate Analysis of the Procrustes coordinates of *Aedes aegypti* by regions. (Abbreviations: CT – Coast, DP – Deccan Peninsula, DT – Desert, HL – Himalaya, NE – North East, SD – Semi Desert)**

Reclassification scores based on populations (*p* < 0.0001) ranged from 88% (Srinagar and Hoshiarpur) to 27% (Visakhapatnam and Srinagar), with an average accuracy of 59%. Srinagar showed the highest average accuracy of 68%, while Raipur exhibited the lowest average reclassification accuracy of 45% (Table 1). This indicates a significant difference in wing shapes among the populations. Whereas, cross-validated reclassification of the six regions yielded significantly high scores (Table 2). Out of 30 comparisons (*p* < 0.0001), only 2 had accuracies below 50% (Himalaya and North East: 46%, Himalaya and Semi Desert: 47%). The Desert region exhibited the highest accuracy (around 70%) compared to other regions. Additionally, the Semi Desert region had the lowest reclassification score, except with the Coast (69%). This implies a significant difference in wing shape among the regions as well.

**Table 1.** Pairwise cross-validated reclassification (%) based on populations. Values below the diagonal correspond to proportion of Group 1 mosquito specimens correctly identified after comparison with Group 2. Values above the diagonal correspond to proportion of Group 2 mosquito specimens correctly identified after comparison with Group 1. (*p* value < 0.0001)

|  |  | Group 2 |  |  |  |  |  |  |  |  |  |  |
| --- | --- | --- | --- | --- | --- | --- | --- | --- | --- | --- | --- | --- |
| Reclassification test |  | GW | HP | IT | JD | KK | KT | NG | RP | SG | SN | VP |
| Group 1 | GW | - | 67 | 53 | 68 | 52 | 38 | 63 | 31 | 85 | 47 | 65 |
|  | HP | 67 | - | 66 | 59 | 81 | 50 | 59 | 54 | 67 | 87 | 60 |
|  | IT | 58 | 67 | - | 64 | 59 | 50 | 67 | 38 | 70 | 47 | 70 |
|  | JD | 88 | 58 | 66 | - | 67 | 50 | 70 | 31 | 78 | 47 | 50 |
|  | KK | 63 | 63 | 75 | 73 | - | 38 | 56 | 54 | 56 | 47 | 60 |
|  | KT | 54 | 63 | 66 | 50 | 59 | - | 52 | 54 | 41 | 53 | 60 |
|  | NG | 46 | 50 | 66 | 73 | 67 | 50 | - | 38 | 59 | 60 | 55 |
|  | RP | 46 | 54 | 56 | 45 | 70 | 63 | 56 | - | 85 | 47 | 60 |
|  | SG | 79 | 63 | 63 | 50 | 59 | 38 | 48 | 69 | - | 87 | 45 |
|  | SN | 63 | 88 | 53 | 59 | 63 | 38 | 74 | 31 | 85 | - | 55 |
|  | VP | 63 | 63 | 81 | 59 | 56 | 50 | 70 | 46 | 52 | 27 | - |

**Table 2.** Pairwise cross-validated species reclassification (%) based on regions. Values below the diagonal correspond to proportion of Group 1 mosquito specimens correctly identified after comparison with Group 2. Values above the diagonal correspond to proportion of Group 2 mosquito specimens correctly identified after comparison with Group 1. (*p* value < 0.0001)

|  |  | **Group 2** |  |  |  |  |  |
| --- | --- | --- | --- | --- | --- | --- | --- |
| **Reclassification test** |  | **CT** | **DP** | **DT** | **HL** | **NE** | **SD** |
| **Group 1** | **CT** | - | 63 | 73 | 68 | 54 | 69 |
|  | **DP** | 66 | - | 71 | 57 | 54 | 50 |
|  | **DT** | 64 | 65 | - | 68 | 79 | 50 |
|  | **HL** | 57 | 65 | 71 | - | 46 | 47 |
|  | **NE** | 68 | 60 | 76 | 68 | - | 59 |
|  | **SD** | 62 | 68 | 65 | 62 | 50 | - |

A significant difference (*p* < 0.0001) was observed in the Mahalanobis distances obtained from pairwise comparisons among the locations ranging from 1.8543 (Itanagar and Guwahati) to 5.2001 (Kota and Hoshiarpur). A similar significant difference was also observed for the region-wise comparison ranging from 1.5969 (Himalaya and Coast) to 2.3511 (Semi Desert and Deccan Peninsula) (Supplementary Tables 2 and 3).

The NJ tree of the population formed two clusters with 100 bootstrap values (Figure 9). Nagpur and Srinagar formed a distinct cluster from the other locations with 100 bootstrap values, while subsequent branching involving all the locations lacked bootstrap support. At the region level (Figure 10), two clusters were formed with 100 bootstrap values. In one cluster, Coast, Deccan Peninsula, and North East grouped together. However, North East region showed more wing shape variation and formed a distinct branch. In the second cluster, Desert, Semi-Desert, and Himalaya regions clustered, with less variation in Desert and Semi Desert regions. However, subsequent branching of both clusters lacked bootstrap support.

**Figure 9.**
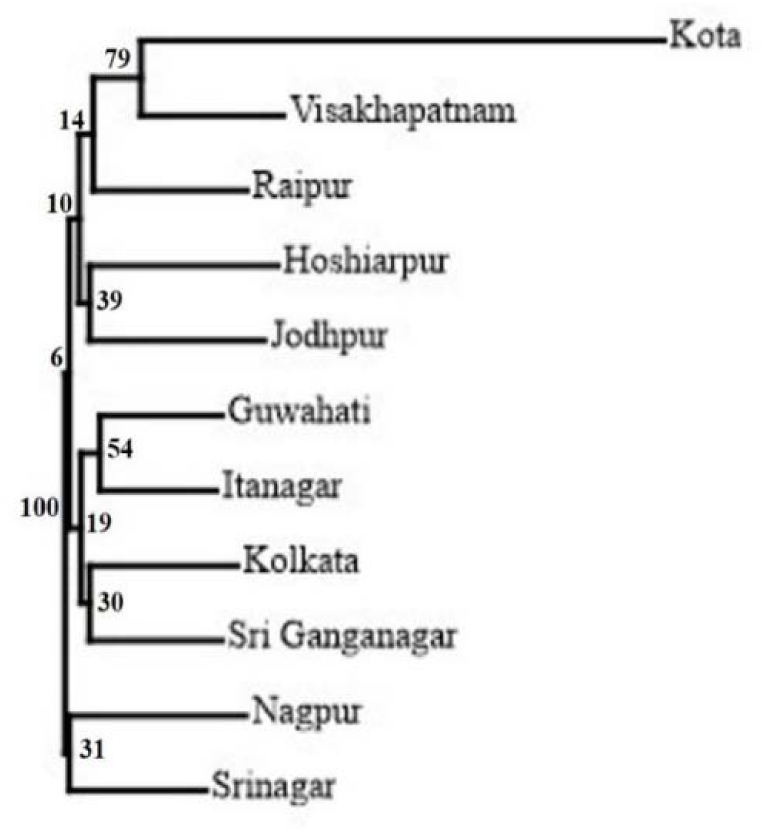
**Neighbor-joining tree based on Mahalanobis distances and computed over 100 bootstrap replicates by populations.**

**Figure 10.**
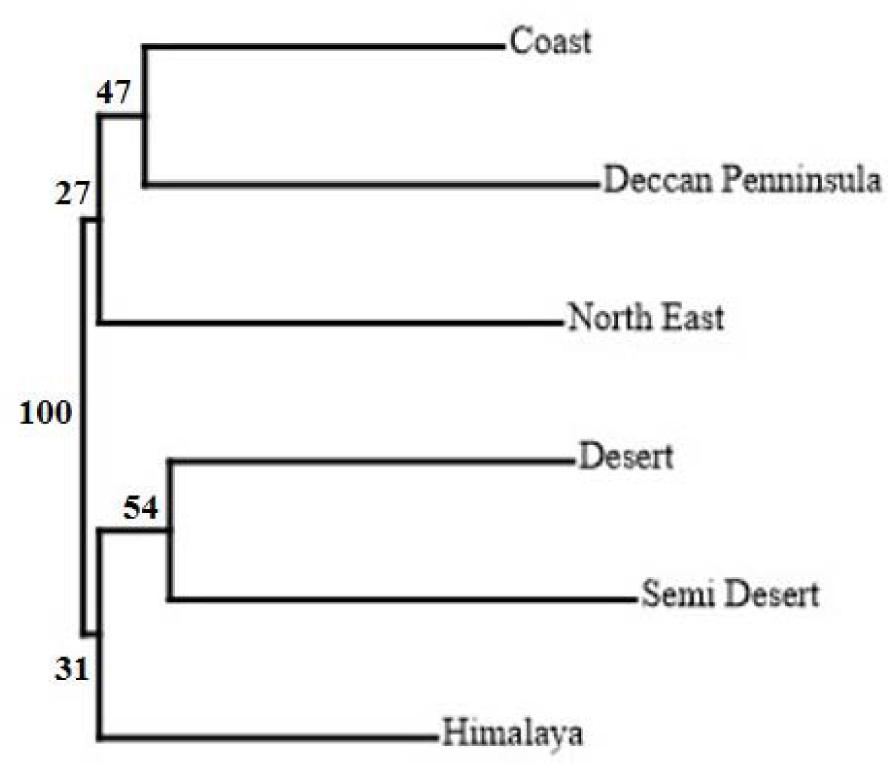
**Neighbor-joining tree based on Mahalanobis distances and computed over 100 bootstrap replicates by regions.**

Multivariate regression of the Procrustes coordinates on log CS shows an allometric effect of wing size on wing shape. (Population: 6.55%, *p* < 0.0001; Region: 2.60%, *p* < 0.0001). We have excluded this effect from the subsequent analysis.

The comparison among the populations in context to their mean Temperature, Relative Humidity, and Precipitation have shown significant differences (*p* < 0.0001) (Supplementary Figure 1). Visakhapatnam (Coast) showed the highest value for temperature (27.36 °C) and RH (76%) whereas Srinagar (9 °C) (Himalaya) and Sri Ganganagar (33.63%) (Desert) showed the lowest values for Temperature and RH respectively. The highest value for precipitation was recorded for Itanagar (5.30 mm) (Himalaya), which is also a North Eastern city, while the lowest was reported for Jodhpur (0.05 mm) (Desert). This indicates that collection locations are statistically significant different in their native environmental conditions.

The multivariate correlation analysis between wing centroid size (CS) and environmental factors did not reveal any significant relationships at the population level. However, at the regional level, only altitude demonstrated a significant positive correlation (82%) with CS (*p* < 0.05). Moreover, temperature exhibited a notable positive relationship of more than 50% with centroid size, both within populations and regions. This information is available in Tables 3 and 4.

**Table 3.** Correlation coefficient matrix of centroid size from 11 Populations of India. (Values below the diagonal represents the corelation coefficient and above the diagonal represents the *p* value)

|  | **Centroid size** | **Latitude** | **Longitude** | **Temperature** | **Relative Humidity** | **Precipitation** | **Altitude** |
| --- | --- | --- | --- | --- | --- | --- | --- |
| **Centroid size** | - | 0.16576 | 0.60279 | 0.082746 | 0.3898 | 0.95151 | 0.2442 |
| **Latitude** | -0.44918 | - | 0.25878 | 0.03828 | 0.43961 | 0.073481 | 0.89846 |
| **Longitude** | -0.17692 | -0.37283 | - | 0.94522 | 0.0077046 | 3.7335E-05 | 0.19489 |
| **Temperature** | 0.54531 | -0.62868 | 0.023546 | - | 0.6408 | 0.78997 | 0.88692 |
| **Relative Humidity** | -0.28838 | -0.26024 | 0.75115 | -0.15887 | - | 0.023331 | 0.26858 |
| **Precipitation** | -<br>0.020838 | -0.55954 | 0.9283 | 0.091086 | 0.67266 | - | 0.37628 |
| **Altitude** | 0.38358 | 0.043711 | -0.42301 | 0.048706 | -0.3658 | -0.29631 | - |

**Table 4.**
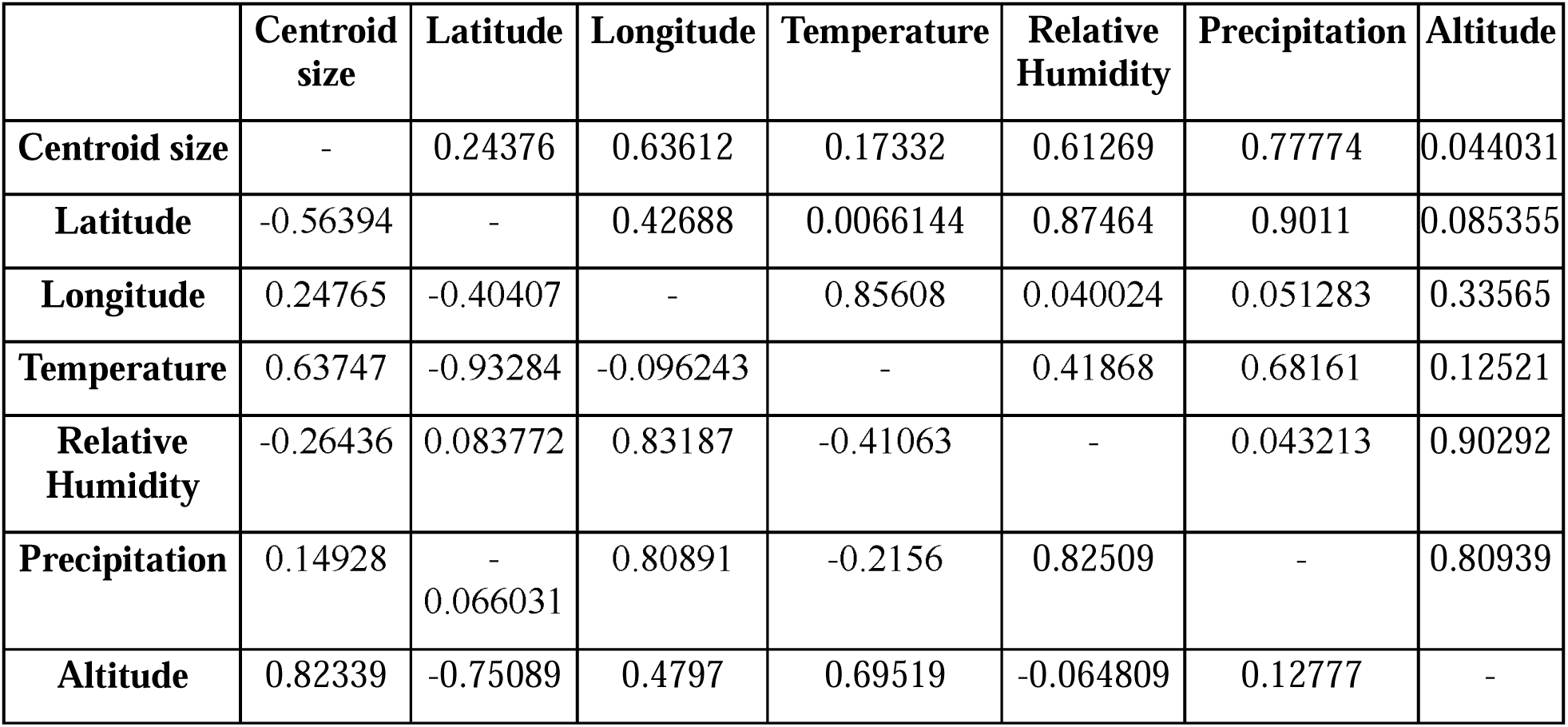
Correlation coefficient matrix of centroid size from six different climatic regions of India. (Values below the diagonal represents the corelation coefficient and above the diagonal represents the *p* value)

## Discussion

*Aedes aegypti* exhibits a remarkable adaptability to anthropogenic modifications, thriving across a diverse range of environmental conditions. To navigate the challenges posed by these dynamic settings, the mosquito must cope with significant selective pressures. Previous research has highlighted the varying impacts of distinct environmental conditions on mosquitoes, influencing their genetics (Sumitha et al., 2023), physiology (Sharma et al., 2023), and morphology (Hounkanrin et al., 2023). Expanding the existing knowledge, the present study analyzed the wing morphometrics of 11 *Ae. aegypti* populations from six geo-climatic regions of India, to understand the influence of native environmental conditions on wing size and shape. Among the 11 *Ae. aegypti* populations, Jodhpur, and Sri Ganganagar populations fall within the Desert region, marked by arid conditions. Kota and Hoshiarpur populations are situated in the Semi-Desert region, characterized by moderately dry climates. Nagpur and Raipur populations represent the Deccan Peninsula, known for its unique landscape features. Srinagar and Itanagar populations are in the Himalayan region, defined by mountainous terrain. Kolkata and Visakhapatnam populations lie along the Coast, experiencing maritime influences. Lastly, Guwahati’s population is positioned in the North East region, distinguished by its specific geographical and climatic attributes. The present investigation has evidenced that these regions and their representative populations are significantly different from one another in context to their climatic conditions including temperature, relative humidity, and precipitation.

Wing size is commonly used as a proxy for overall mosquito size (Dujardin, 2008), and it can influence various physiological and behavioral aspects of mosquitoes (Yeap et al., 2013). Larger mosquitoes, characterized by greater wing size, often exhibit distinct metabolic profiles, enhanced blood-feeding capacities, and improved flight capabilities compared to their smaller size mosquitoes (Yeap et al., 2013). These factors can collectively affect the mosquito’s ability to transmit diseases. For example, larger mosquitoes are less susceptible to viral infections, potentially influencing the dynamics of virus transmission within mosquito populations (Alto et al., 2008). Therefore, wing morphometric studies have practical implications for formulating more effective vector management strategies in specific areas, emphasizing the importance of considering local environmental conditions in such interventions. Our results, derived from centroid size, indicate that the Deccan Peninsula region (4.46 ± 0.10 mm) supports larger wing size mosquitoes, represented by Nagpur (4.48 ± 0.06 mm) and Raipur (4.40 mm ± 0.09) populations. In contrast, the Himalayan region (2.04 ± 0.09 mm) recorded smaller wing sizes, as evidenced by measurements from Srinagar (1.92 ± 0.08 mm) and Itanagar (2.09 ± 0.05 mm). This observed phenomenon can be attributed to the specific environmental conditions in the Deccan Peninsula, potentially providing favorable conditions for the development of larger mosquito wings. Conversely, the Himalayan region, characterized by adverse environmental conditions, particularly lower temperatures, may interfere with mosquito development, resulting in shorter wing lengths. Our correlation analysis of centroid size with temperature and altitude further supports this observation.

While wing size provides valuable information, it is acknowledged that it is more susceptible to environmental changes (Lorenz et al., 2017). In Thailand, the wing size of *Ae. albopictus* populations appear to be influenced by climatic conditions, whereas wing shape can reveal heritable intraspecific and regional differences (Vargas et al., 2013). Therefore, caution should be exercised when interpreting wing size data. In contrast, wing shape, as highlighted by research, proves to be more resilient to environmental variations and can predict population structure in certain species (Carvajal et al., 2016, Rodríguez-Zabala et al., 2016, Krtinić et al., 2016). The morphospace analysis, conducted through Canonical Variate Analysis (CVA), reveals statistically significant differences in wing forms among populations and regions. The Semi Desert region (Kota population) contributed a larger morphospace than the other regions, revealing a greater variety of wing shapes. The cross-validated reclassification results also show that wing shape varies significantly between regions. Only two of the 30 comparisons had accuracies less than 50% (Himalaya and North East (46%), Himalaya and Semi Desert (47%)). Similarly, for populations, an average accuracy of 59% was obtained from 110 comparisons. This suggests that these populations constitute a part of a micro-geographic region, which contributes to significant population structure in *Ae. aegypti*, driven by variable climate conditions.

The NJ tree produced in the present study also supports the results of the wing shape CVA and reclassification test. However, the study observes that populations from the same region did not consistently form the same cluster, suggesting the absence of a directional trend in wing shape characteristics among populations and regions.

## Conclusion

The current study highlights the variation in wing size and shape of *Ae. aegypti* sampled from different geo-climatic regions of India. The analysis of the wing size attribute illustrates that populations from colder regions (Himalaya) have smaller wing sizes, whereas tropical regions with wet and dry conditions (Deccan Peninsula) have larger mosquito wings. We also identified significant variations in wing shape among populations and regions. However, we did not find a particular directional trend for this characteristic. This shows that numerous environmental factors influence wing shape differences. The outcomes of this study will help in understanding the emerging population structure of *Ae. aegypti* in India. These insights may serve as a foundation for more targeted and effective strategies in the surveillance and management of this vector species.

## Acknowledgement

The authors thanks to Dr. Surya Singh, Scientist-B, ICMR-National institute for Research in Environmental Health, Bhopal, India, for providing technical support.

## Author Contributions

GS: Conceptualization, Data Curation, Resources, Software, Validation, Writing – original draft. RB: Supervision, Resources, Software, Writing - review & editing. DKS: Conceptualization, Supervision, Funding acquisition, Project administration, Writing – original draft.

## Ethical statement

The study does not require any ethical approval.

## Supplementary Figures

Supplementary Figure 1. Comparison among the populations of *Aedes aegypti* concerning their mean Temperature, Relative Humidity, and Precipitation.

## Supplementary Tables

Supplementary Table 1. Details of female *Aedes aegypti* collection sites with their mean centroid sizes.

Supplementary Table 2. Pairwise Mahalanobis distances based on populations.

Supplementary Table 3. Pairwise Mahalanobis distances based on climatic regions.

## Supplementary File

Supplementary File 1. Details of raw wing coordinates of each *Ae. aegypti* specimen.

