## Supplementary for "Effect of environmental conditions on the wing morphometric variation in *Aedes aegypti* (Diptera: Culicidae) in India": Supplementary Figure_27012024.docx

**Supplementary Figure 1. Comparison among the populations of *Aedes aegypti* concerning their mean Temperature, Relative Humidity, and Precipitation.**

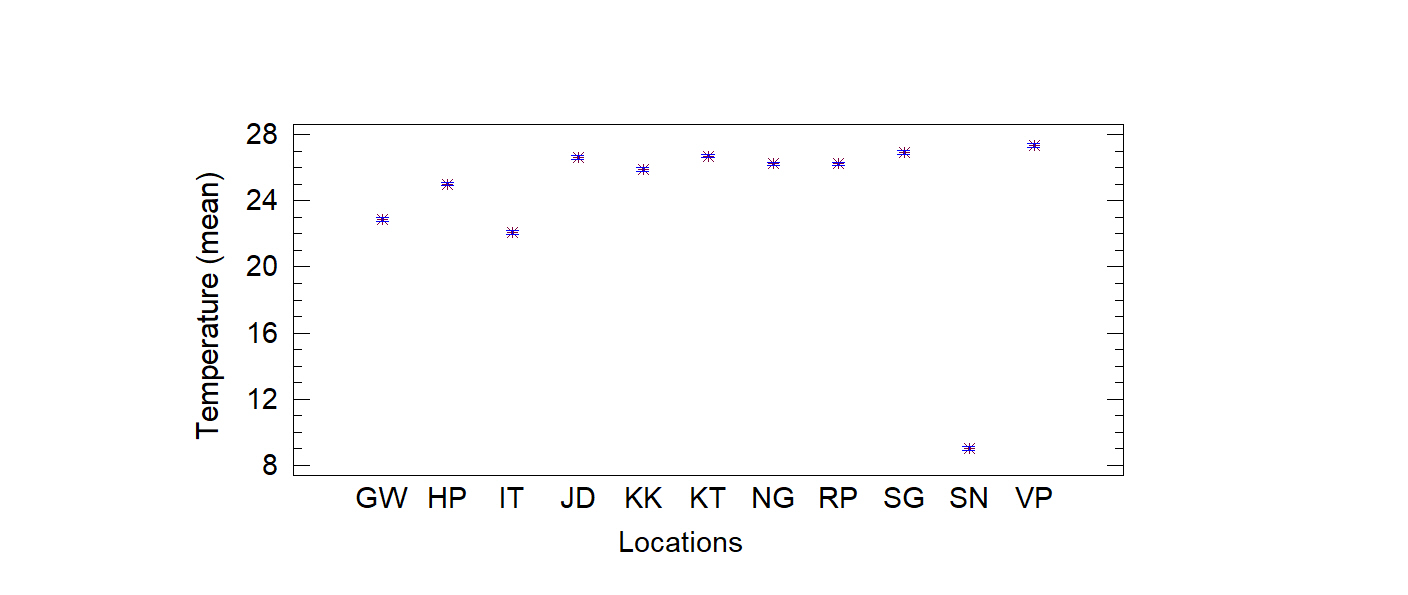

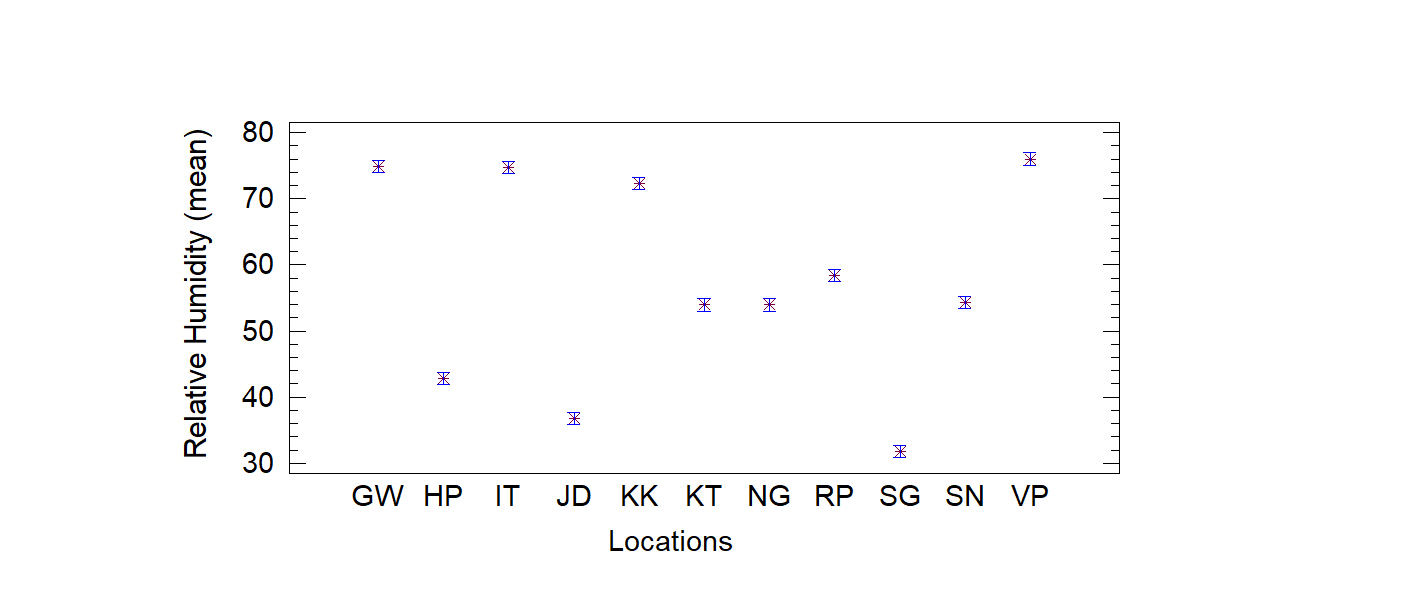

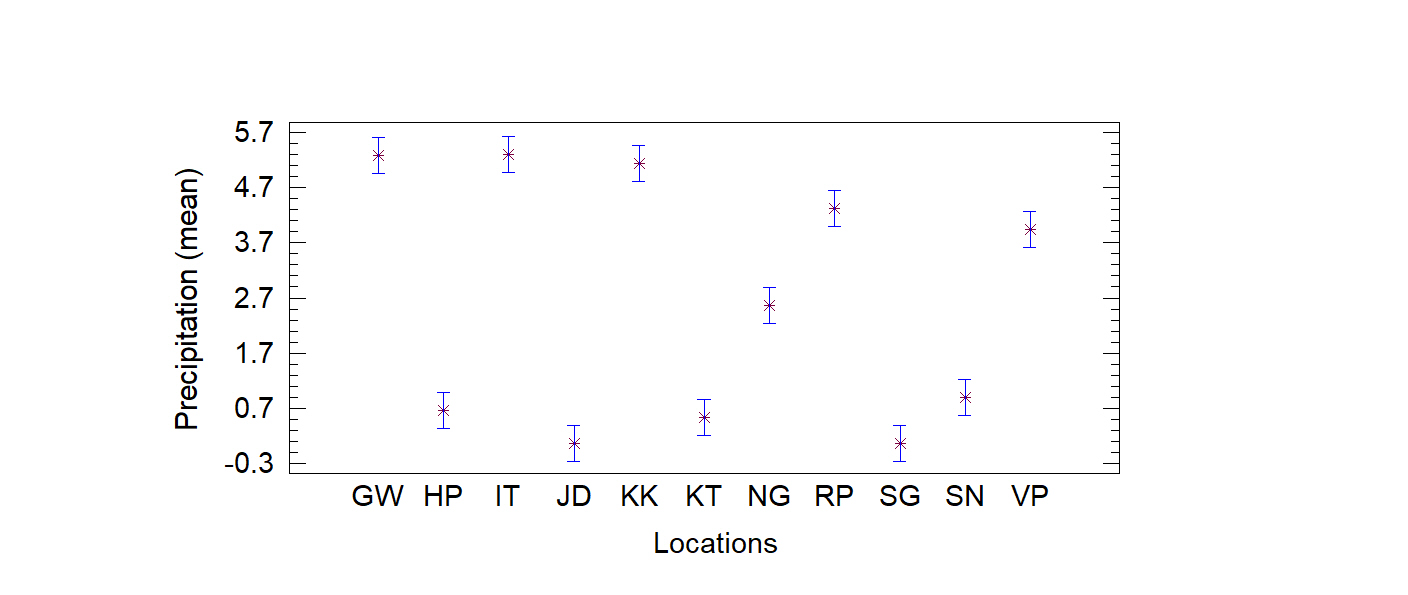
