## Supplementary for "Effect of environmental conditions on the wing morphometric variation in *Aedes aegypti* (Diptera: Culicidae) in India": Supplementary Table_27012024.docx

**Table 1. Details of female *Aedes aegypti* collection sites with their mean centroid sizes.**

| Region | Location | Sites | Latitude | Longitude | No. of female specimen | Mean centroid size (mm) |
| --- | --- | --- | --- | --- | --- | --- |
| Himalaya | Srinagar | SN-01 | 34°01'56.2"N | 74°49'19.7"E | 12 | 1.947593214 |
|  |  | SN-02 | 34°01'56.2"N | 74°49'19.7"E | 3 | 1.848982424 |
|  | Itanagar | IT_01 | 27°05'52.1"N | 93°38'05.6"E | 13 | 2.012967325 |
|  |  | IT_02 | 27°06'01.1"N | 93°37'56.6"E | 11 | 2.094475588 |
|  |  | IT_03 | 27°05'42.4"N | 93°37'12.6"E | 8 | 2.221814659 |
| Desert | Jodhpur | JD_01 | 26°17'45.1"N | 73°01'23.0"E | 2 | 3.86578107 |
|  |  | JD_02 | 26°16'40.7"N | 73°01'15.3"E | 4 | 3.110169335 |
|  |  | JD_03 | 26°16'10.9"N | 73°00'53.6"E | 4 | 3.639543788 |
|  |  | JD_04 | 26°16'08.9"N | 72°59'53.3"E | 4 | 3.513478926 |
|  |  | JD_05 | 26°16'26.0"N | 73°01'50.5"E | 4 | 3.819760336 |
|  |  | JD_06 | 26°14'49.1"N | 72°58'06.1"E | 3 | 3.615287704 |
|  |  | JD_07 | 26°15'45.1"N | 72°58'04.5"E | 1 | 3.179182993 |
|  | Sri Ganganagar | SG_01 | 29°55'55.5"N | 73°52'22.1"E | 6 | 4.685994402 |
|  |  | SG_02 | 29°55'20.3"N | 73°52'04.5"E | 2 | 4.076342951 |
|  |  | SG_03 | 29°55'29.0"N | 73°52'19.7"E | 7 | 4.446867603 |
|  |  | SG_04 | 29°55'33.0"N | 73°52'35.6"E | 7 | 3.608397105 |
|  |  | SG_05 | 29°55'37.1"N | 73°52'39.7"E | 3 | 4.445714954 |
|  |  | SG_06 | 29°55'36.2"N | 73°52'37.4"E | 2 | 4.862498354 |
| Semi-desert | Kota | KT_01 | 25°07'39.0"N | 75°49'32.9"E | 3 | 4.659231471 |
|  |  | KT_02 | 25°12'19.1"N | 75°50'09.6"E | 2 | 3.590012043 |
|  |  | KT_03 | 25°10'41.9"N | 75°53'06.0"E | 3 | 2.838199324 |
|  | Hoshiarpur | HP_01 | 31°31'13.1"N | 75°54'44.5"E | 15 | 2.301833237 |
|  |  | HP_02 | 31°31'27.7"N | 75°55'14.6"E | 5 | 2.261132794 |
|  |  | HP_03 | 31°31'32.9"N | 75°55'02.8"E | 4 | 2.476762277 |
| Deccan Peninsula | Nagpur | NG_01 | 21°09'04.2"N | 79°06'07.3"E | 4 | 4.316189692 |
|  |  | NG_02 | 21°09'04.1"N | 79°05'39.5"E | 6 | 4.156105336 |
|  |  | NG_03 | 21°07'36.2"N | 79°06'09.5"E | 3 | 4.730758283 |
|  |  | NG_04 | 21°08'17.0"N | 79°06'58.0"E | 3 | 4.484089562 |
|  |  | NG_05 | 21°09'13.0"N | 79°05'37.3"E | 5 | 4.446609638 |
|  |  | NG_06 | 21°08'32.9"N | 79°02'47.1"E | 2 | 4.653837784 |
|  |  | NG_07 | 21°09'08.3"N | 79°05'17.5"E | 4 | 4.960372855 |
|  | Raipur | RP_01 | 21°13'18.6"N | 81°36'50.8"E | 1 | 4.002799971 |
|  |  | RP_02 | 21°15'47.6"N | 81°37'16.8"E | 1 | 4.552695789 |
|  |  | RP_03 | 21°15'48.9"N | 81°38'25.9"E | 1 | 4.926744728 |
|  |  | RP_04 | 21°14'51.0"N | 81°38'13.5"E | 2 | 4.657139693 |
|  |  | RP_05 | 21°12'08.2"N | 81°38'13.5"E | 1 | 4.002158862 |
|  |  | RP_06 | 21°17'36.3"N | 81°36'59.1"E | 2 | 4.477564053 |
|  |  | RP_07 | 21°17'39.0"N | 81°38'39.5"E | 1 | 3.586235925 |
|  |  | RP_08 | 21°18'23.2"N | 81°39'07.8"E | 2 | 4.284846943 |
|  |  | RP_09 | 21°17'18.9"N | 81°40'10.1"E | 2 | 4.676153927 |
| Coast | Kolkata | KK_01 | 22°34'00.8"N | 88°21'28.7"E | 3 | 2.055730014 |
|  |  | KK_02 | 22°33'45.9"N | 88°21'21.8"E | 11 | 1.995915395 |
|  |  | KK_03 | 22°34'53.7"N | 88°19'54.0"E | 8 | 1.968650564 |
|  |  | KK_04 | 22°34'48.0"N | 88°17'19.0"E | 5 | 2.037860064 |
|  | Visakhapatnam | VP_01 | 17°43'34.4"N | 83°18'35.2"E | 3 | 4.20767539 |
|  |  | VP_02 | 17°44'48.9"N | 83°19'39.8"E | 2 | 4.153016052 |
|  |  | VP_03 | 17°44'05.1"N | 83°17'57.3"E | 1 | 3.882927175 |
|  |  | VP_04 | 17°44'34.8"N | 83°17'51.0"E | 2 | 3.982702655 |
|  |  | VP_05 | 17°44'42.2"N | 83°18'48.2"E | 2 | 4.231631437 |
|  |  | VP_06 | 17°43'58.4"N | 83°19'42.9"E | 2 | 4.23193058 |
|  |  | VP_07 | 17°43'57.1"N | 83°20'21.5"E | 1 | 4.048139147 |
|  |  | VP_08 | 17°45'48.4"N | 83°20'01.8"E | 1 | 4.184224128 |
|  |  | VP_09 | 17°48'14.1"N | 83°21'39.1"E | 1 | 3.98954421 |
|  |  | VP_10 | 17°48'55.4"N | 83°22'05.4"E | 2 | 3.510658972 |
|  |  | VP_11 | 17°49'13.9"N | 83°21'39.4"E | 2 | 4.193526272 |
|  |  | VP_12 | 17°49'32.3"N | 83°21'29.3"E | 1 | 4.425789829 |
| North-east | Guwahati | GW_01 | 26°10'02.9"N | 91°44'48.2"E | 8 | 4.298166038 |
|  |  | GW_02 | 26°10'08.0"N | 91°45'33.5"E | 9 | 4.435266055 |
|  |  | GW_03 | 26°09'53.8"N | 91°41'30.7"E | 4 | 3.973882779 |
|  |  | GW_04 | 26°09'51.8"N | 91°41'24.0"E | 1 | 4.558775762 |
|  |  | GW_05 | 26°10'23.8"N | 91°44'54.7"E | 2 | 5.160488768 |
| Total |  |  |  |  | **239** |  |

**Table 2. Pairwise Mahalanobis distances based on populations**

|  | **Guwahati** | **Hoshiarpur** | **Itanagar** | **Jodhpur** | **Kolkata** | **Kota** | **Nagpur** | **Raipur** | **Sri Ganganagar** | **Srinagar** | **Visakhapatnam** |
| --- | --- | --- | --- | --- | --- | --- | --- | --- | --- | --- | --- |
| **Guwahati** | - | 2.3848 | 1.8543 | 2.8577 | 2.24 | 4.355 | 2.4922 | 2.4197 | 2.3613 | 2.2271 | 2.3751 |
| **Hoshiarpur** | 2.3848 | - | 2.4794 | 2.7827 | 3.1684 | 5.2001 | 3.1592 | 2.7688 | 2.6222 | 2.4908 | 3.2097 |
| **Itanagar** | 1.8543 | 2.4794 | - | 2.5555 | 2.01 | 4.2685 | 2.5986 | 2.2207 | 2.2433 | 2.1573 | 2.7889 |
| **Jodhpur** | 2.8577 | 2.7827 | 2.5555 | - | 2.797 | 4.2792 | 2.839 | 2.7106 | 2.6648 | 2.5296 | 3.4192 |
| **Kolkata** | 2.24 | 3.1684 | 2.01 | 2.797 | - | 4.3792 | 2.6651 | 2.6191 | 2.1788 | 2.4142 | 2.9656 |
| **Kota** | 4.355 | 5.2001 | 4.2685 | 4.2792 | 4.3792 | - | 4.9204 | 4.3915 | 4.3912 | 4.1449 | 3.9108 |
| **Nagpur** | 2.4922 | 3.1592 | 2.5986 | 2.839 | 2.6651 | 4.9204 | - | 2.8837 | 2.6636 | 2.3317 | 2.7838 |
| **Raipur** | 2.4197 | 2.7688 | 2.2207 | 2.7106 | 2.6191 | 4.3915 | 2.8837 | - | 2.284 | 2.7354 | 2.4427 |
| **Sri Ganganagar** | 2.3613 | 2.6222 | 2.2433 | 2.6648 | 2.1788 | 4.3912 | 2.6636 | 2.284 | - | 2.3269 | 2.8187 |
| **Srinagar** | 2.2271 | 2.4908 | 2.1573 | 2.5296 | 2.4142 | 4.1449 | 2.3317 | 2.7354 | 2.3269 | - | 2.5674 |
| **Visakhapatnam** | 2.3751 | 3.2097 | 2.7889 | 3.4192 | 2.9656 | 3.9108 | 2.7838 | 2.4427 | 2.8187 | 2.5674 | - |

**Table 3. Pairwise Mahalanobis distances based on climatic regions**

|  | **Coast** | **Deccan Peninsula** | **Desert** | **Himalaya** | **North East** | **Semi Desert** |
| --- | --- | --- | --- | --- | --- | --- |
| **Coast** | - | 1.6958 | 1.8996 | 1.5969 | 1.7433 | 2.1329 |
| **Deccan Peninsula** | 1.6958 | - | 1.8231 | 1.7863 | 2.0617 | 2.3511 |
| **Desert** | 1.8996 | 1.8231 | - | 1.7359 | 2.2225 | 1.79 |
| **Himalaya** | 1.5969 | 1.7863 | 1.7359 | - | 1.7054 | 1.6933 |
| **North East** | 1.7433 | 2.0617 | 2.2225 | 1.7054 | - | 1.9685 |
| **Semi Desert** | 2.1329 | 2.3511 | 1.79 | 1.6933 | 1.9685 | - |
